# Multi-Class Classification of Cannabis and Alcohol Use Disorder: Identifying Common and Substance-Specific Neural Circuits

**DOI:** 10.1101/2025.10.24.684424

**Authors:** Ayaka Kato, Kaustubh R. Kulkarni, Nikash R. Das, Matthew Heflin, Vincenzo G. Fiore, Francesca M Filbey, Xiaosi Gu

**Author notes:** These authors contributed equally (Ayaka Kato, Kaustubh R. Kulkarni).

## Abstract

Machine learning approaches have advanced the identification of neural signatures of substance use, particularly through case-control comparisons and network-level analyses. However, most studies have focused on single substances in isolation, making it difficult to directly compare shared and distinct network computations and limiting their generalizability, given that many individuals engage in polysubstance use. Here, we introduce an explainable, connectivity-based multiclass classification framework that models distributed brain network organization to delineate both shared and substance-specific mechanisms of addiction. This design enables direct comparison among cannabis users, alcohol users, and controls, revealing network computations that uniquely differentiate each group. Using functional connectivity features from cue-induced craving tasks, we classified cannabis users (n=166), alcohol users (n=101), and healthy controls (n=238), achieving high out-of-sample accuracy (87% for cannabis, 69% for alcohol, and 73% for controls). Functional connectivity-based models consistently outperformed activation-based models, highlighting the importance of inter-regional network properties for biomarker development. Network analysis further revealed dorsal prefrontal, cingulate, and precuneus hubs consistent with meta-analytic craving networks and substance-specific connectivity motifs enhanced prefrontal coupling in cannabis users and greater insular-striatal integration in alcohol users. These results demonstrate that distinct network configurations define different substance-use profiles, advancing interpretable biomarkers for addiction neuroscience.

## Main

Substance use disorders (SUDs), including cannabis and alcohol use disorder, are prevalent, chronic conditions that contribute to significant global disease burden and cognitive dysfunction. Despite extensive neuroimaging research implicating aberrant reward, salience, and executive control circuits in SUDs (Ekhtiari et al., 2024), the development of reliable, interpretable, and clinically useful biomarkers remains limited. While diagnostic criteria for substance use disorders are well established based on symptomatology, current neurobiological assessments remain limited. Most objective measures rely on detecting psychoactive substances or their metabolites in biological samples, which do not reflect the underlying network-level dysfunction. Furthermore, existing neuroimaging models often simplify circuits shared across substances, hindering the identification of substance-specific mechanisms. Addressing these limitations is critical for bridging neurobiological insights and clinical translation in addiction medicine.

Functional magnetic resonance imaging (fMRI) has become the dominant modality for investigating brain function in SUDs, with over 250 clinical trials incorporating it as an outcome measure, most of which involve cue exposure paradigms, as reviewed by Ekhtiari et al. (2024). Many studies have adopted activity-based models that derive biomarkers from localized, task-evoked activations to investigate the neural mechanisms of addiction. A notable advance in this domain is the Neurobiological Craving Signature (NCS) introduced by Koban et al. (2023a), which applied machine learning to distributed activity patterns to predict craving across substances. While this represents a significant methodological step forward, such models still focus primarily on voxel-wise activations and may not adequately reflect the distributed, systems-level pathology underlying addiction.

To capture the network-level dysfunctions characteristic of SUDs—such as impaired cognitive control, aberrant salience processing, and heightened cue reactivity—connectivity-based models offer a compelling alternative. These approaches quantify statistical dependencies among brain regions, providing insight into the brain’s intrinsic network organization. Prior research has shown that connectivity-based features generalize more effectively across individuals and yield biologically meaningful insights into addiction-related circuits (Goldstein & Volkow, 2011; Zilverstand et al., 2018; Sutherland et al., 2012). In our previous work, we demonstrated that connectivity-based classifiers outperform activity-based models in identifying chronic cannabis users, offering both improved predictive accuracy and interpretability (Kulkarni et al., 2022). However, comparative evaluations across different SUD populations remain limited.

In the present study, we expand this framework to both cannabis and alcohol use disorders, providing one of the first systematic comparisons of activity- and connectivity-based classification models across multiple substance use populations. We also introduce a three-way classification paradigm that distinguishes cannabis users, alcohol users, and healthy controls—mirroring the diagnostic complexity encountered in real-world clinical contexts. Although multi-class prediction poses methodological challenges (Arbabshirani et al., 2017), it is essential for developing translational tools that extend beyond conventional binary case–control designs (Dinga et al., 2021).

Our findings reveal that connectivity-based models consistently outperform activity-based models across binary, cross-substance, and three-way classification tasks. Moreover, these models identify both substance-specific and shared connectivity signatures involving key hubs in the prefrontal, cingulate, and insular regions—areas previously implicated in craving and self-regulation. These results converge with meta-analytic findings from the Neurosynth database, underscoring the clinical relevance of large-scale network disruptions in addiction.

In summary, we present one of the first cross-substance, multi-class evaluations of brain-based classification models in SUDs. By combining machine learning with functional connectivity features derived from cue-induced craving tasks, our approach achieves both high predictive performance and biological interpretability. These results support the utility of network-informed models as clinically actionable neuromarkers that can inform future efforts toward precision psychiatry in addiction.

## Results

### Participant Characteristics and Classification Framework

We first established the participant characteristics and validated a unified classification framework to test whether connectivity-based features improve classification accuracy relative to activity-based features. Participants included 166 cannabis users (mean age = 30.1 years; 60% male), 136 alcohol users (mean age = 28.3 years; 77% male), and two independent groups of healthy control (HC) subjects: 124 controls matched to the cannabis users (mean age = 27.4 years; 43% male) and 132 controls matched to the alcohol users (mean age = 29.75 years; 48% male). Cannabis and alcohol use severity measures, including past 90-day use frequency and Alcohol Use Disorders Identification Test (AUDIT) scores, are summarized in Supplementary Table 1. The cannabis user dataset was derived from two standardized cue-reactivity studies (Filbey et al., 2009; Filbey et al., 2016), and the alcohol user dataset from an alcohol-taste cue-reactivity study (Filbey et al., 2008).

To address our primary questions—(1) whether functional connectivity features enable more accurate classification than activity-based features, and (2) which brain networks contribute to group differentiation—we developed an analysis pipeline combining feature extraction, model training, and network-level interpretation (Fig. 1). For each participant, we derived voxel-wise activity maps and inter-regional functional connectivity matrices from task-based fMRI data. These were used as inputs to four types of linear classifiers: L1- and L2-penalized logistic regression, and linear support vector classifiers (SGDClassifier variants). Classification performance was assessed using cross-validation on training data and accuracy/recall on independent test sets.

**Figure 1.**
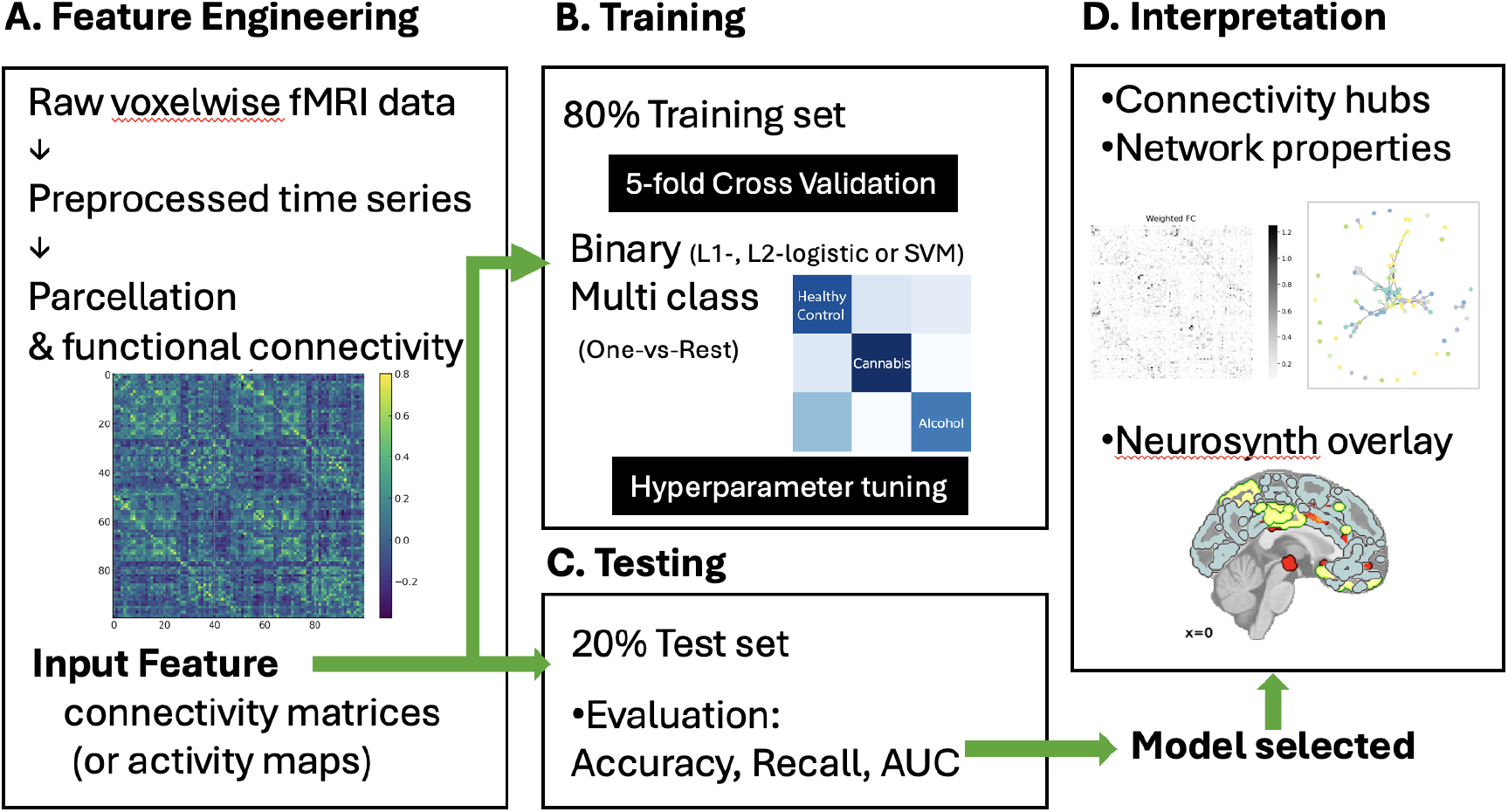
Machine learning pipeline for multi-class classification of substance use. (A) Preprocessed functional MRI (fMRI) time series from a cue-induced craving task were used to derive two types of features: voxel-wise activity maps and functional connectivity matrices. Activity maps were z-scored contrast images, while connectivity matrices were computed using pairwise Pearson correlations between 90 functional regions of interest defined by the Stanford functional atlas. (B) Feature sets were divided into a training set (80% of participants) and a held-out testing set (20%). The training set was further split into 5 folds for cross-validation (CV). For each fold, 4 folds were used for training (blue) and 1 fold for validation (green). Four linear classification algorithms were evaluated: L1- and L2-penalized logistic regression and L1- and L2-penalized linear support vector classification (SVC). Hyperparameters, including regularization strength, were optimized based on cross-validated accuracy, area under the curve (AUC), and precision/recall metrics. (C) The best-performing model was retrained on the full training set and evaluated on the independent testing set to assess generalization. Multi-class classification was implemented using a One-vs-Rest (OvR) strategy to differentiate cannabis users, alcohol users, and healthy controls. Classification performance was summarized using confusion matrices and recall scores for each group. (D) Subject-specific connectivity matrices were further analyzed by weighting connections with model-derived coefficients. Network analyses, including degree centrality and community structure, were performed to identify brain hubs critical for substance use classification. Identified hubs were compared to meta-analytic craving-related brain maps derived from Neurosynth for biological validation. AUC, area under the curve; CV, cross-validation; OvR, One-vs-Rest; fMRI, functional magnetic resonance imaging.

**Figure 2.**
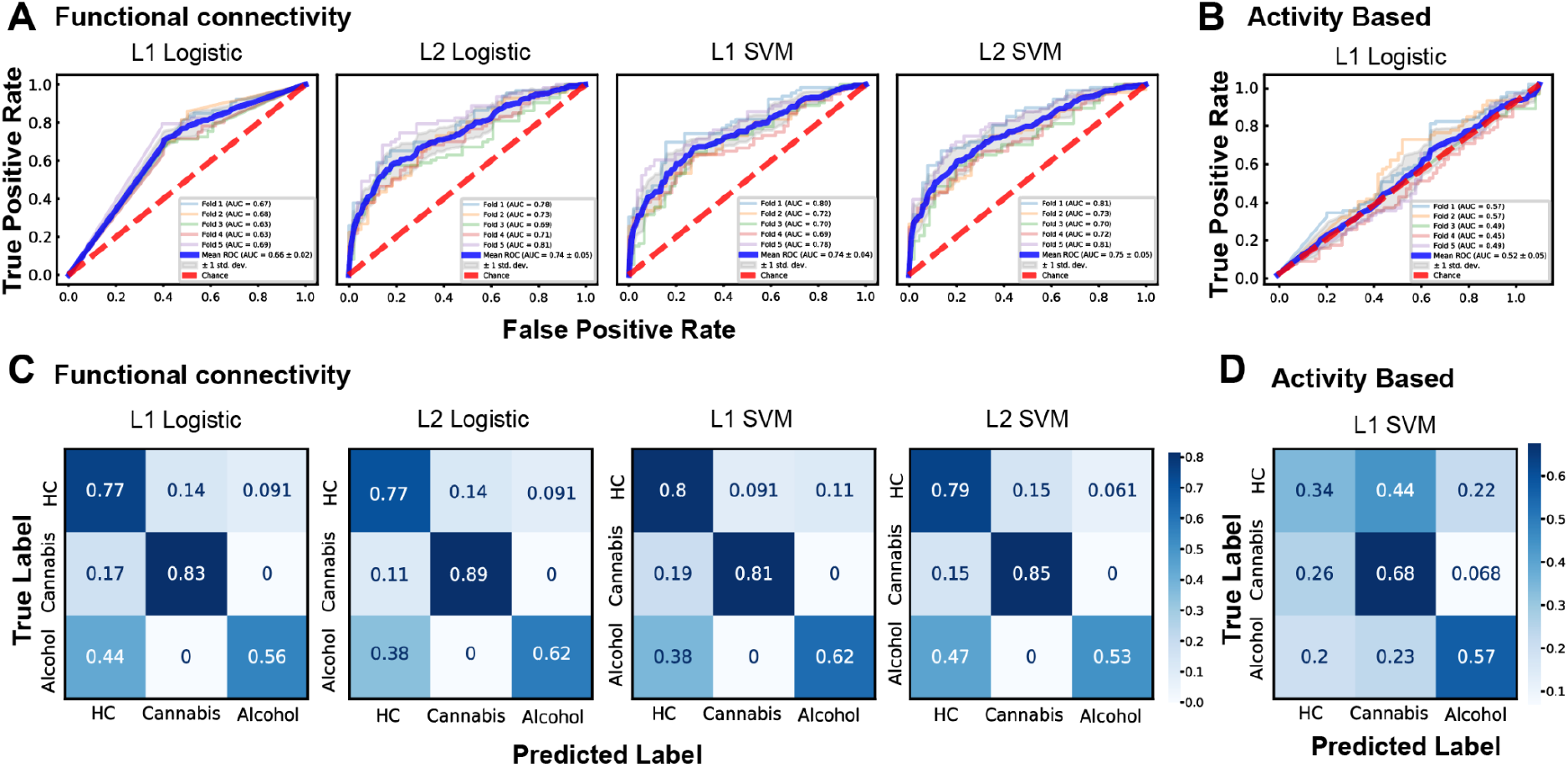
Classification Performance Across Binary and Multiclass Tasks. (A) Receiver operating characteristic (ROC) curves for four connectivity-based classifiers (L1-penalized logistic regression, L2-penalized logistic regression, L1-penalized linear support vector machine, and L2-penalized linear support vector machine) trained on the aggregated binary classification task (healthy controls vs. substance users). (B) Receiver operating characteristic (ROC) curves for four activation-based classifiers trained on the same aggregated binary classification task. (C) Confusion matrices for the four connectivity-based classifiers evaluated on the three-way classification task (healthy controls, cannabis users, alcohol users). (D) Confusion matrix for the best-performing activation-based classifier (SGD classifier with L1 regularization) evaluated on the three-way classification task. ROC curves represent mean performance across cross-validation folds. Confusion matrices display model predictions on the independent test set.

### Functional Connectivity Enables Robust and Balanced Classification of Substance Use Groups

To evaluate the neural separability of individuals with substance use from healthy controls (HC), we first conducted binary classification analyses using both functional connectivity and activation-based features from task-based fMRI. Connectivity-based classifiers demonstrated reliable performance across both alcohol and cannabis groups. For alcohol users versus HC, cross-validated models achieved mean accuracy up to 58.7% (L2-LinearSVC, AUC=0.6068), with a held-out test accuracy of 67.2%. For cannabis users, classification performance was notably higher, with cross-validated accuracy reaching 77.0% (L2-LinearSVC, AUC=0.8123), and a held-out test accuracy of 80.7%. In contrast, activation-based models performed consistently worse. For alcohol users, CV accuracy remained near chance (max 46.9%), with test accuracy under 51%, and AUCs hovering around 0.48–0.49. Cannabis classification showed similar limitations, with mean CV accuracy around 60% only for L2-penalized models (test accuracy also ∼62.7%), while recall for HC remained poor. These results indicate that functional connectivity features are more robust and discriminative for substance use classification than voxel-level task activation maps.

Next, to investigate shared versus substance-specific neural patterns, we trained a cross-substance model: classifiers trained on cannabis users were tested on alcohol users and vice versa. Connectivity-based models yielded above-chance accuracy in both directions, especially when training on alcohol and testing on cannabis (AUC up to 0.548, F1 = 0.703, L2-LinearSVC). However, models trained on cannabis and tested on alcohol performed less well (AUC = 0.602, F1 = 0.550, L2-LinearSVC), suggesting asymmetry in feature generalizability. Activation-based models, meanwhile, failed to generalize across substance groups in both directions, with cross-evaluation accuracies remaining between 50–58%, and F1-scores well below 0.61, further supporting the limited transferability of activation features.

To assess commonalities across substance use conditions, we aggregated both substance groups and performed binary classification against HC. The connectivity-based aggregated model reached a mean CV accuracy of 67.7% (L2-LinearSVC), with test accuracy of 73.7%, suggesting partially shared but discriminable connectivity features. In contrast, the activation-based aggregated model plateaued at CV accuracy of 56.8%, and test accuracy of 58.8%, underscoring its limited generalizability across groups.

Finally, we implemented multi-class classification using a One-vs-Rest (OvR) framework to simultaneously distinguish healthy controls (HC), cannabis users, and alcohol users. Among the connectivity-based models, L2-penalized logistic regression achieved the best overall performance, with a mean cross-validation accuracy of 80.3% and a held-out test accuracy of 78.3%. Class-wise recall reached 89% for cannabis, 62% for alcohol, and 77% for HC, reflecting both high precision and balanced sensitivity across substance groups. These results highlight the model’s ability to capture distributed connectivity patterns that generalize well across individuals with different substance use profiles. In contrast, the activation-based OvR model achieved only 52% test accuracy, with class-wise recall of 68% for cannabis, 57% for alcohol, and just 34% for healthy controls. Misclassifications were broadly distributed across classes, with no consistent skew toward a particular group. These results suggest that while voxel-level activation features may capture some group differences—particularly among cannabis users—they fail to support balanced and generalizable multi-class classification. Overall, these findings emphasize the superiority of connectivity-based features in capturing distributed neural signatures that enable robust differentiation among complex psychiatric populations.

### Network-level substrates of predictive classification features

To characterize the large-scale architecture underlying predictive classification, we examined spatial and network-level distributions of degree centrality (DC) derived from coefficient-weighted connectivity matrices of the best-performing one-vs-rest models (Fig. 3). Spatial maps revealed distributed but convergent configurations of predictive hubs across groups, many of which overlapped with the *drug*-related meta-analytic map from Neurosynth (FDR q < 0.01). For each group, the top 20 regions were selected based on the highest mean degree centrality values across cross-validated folds of the OVR_L2_logistic classifier, ensuring consistency in hub identification across models.

**Figure 3.**
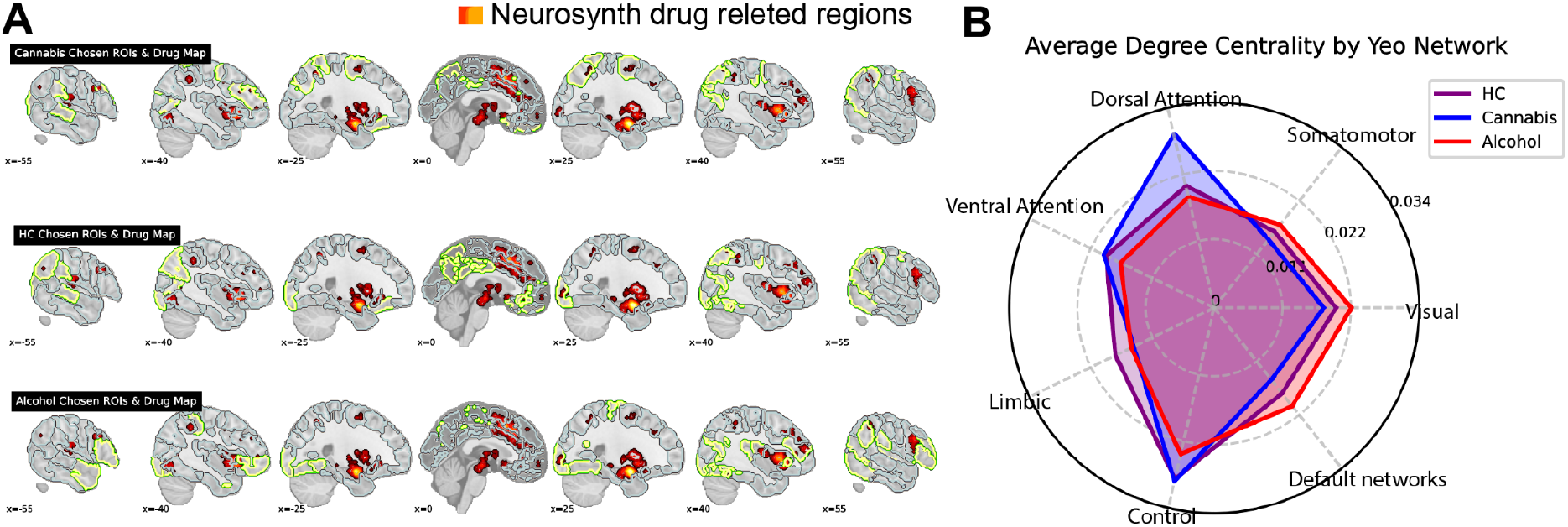
Degree centrality distributions of predictive brain parcels across groups. (A) Brain maps display the top 20 regions with the highest mean degree centrality (DC) from the best-performing one-vs-rest classifier (OVR_L2_logistic). These predictive hubs are outlined in yellow–green and overlaid with the *drug*-related meta-analytic map from Neurosynth (uniformity test, FDR q < 0.01; red gradient). Overlapping regions appear prominently in the medial prefrontal cortex (mPFC), posterior cingulate cortex (PCC), orbitofrontal cortex (OFC), and insula, consistent with canonical reward and salience systems. Group-specific spatial patterns highlight substance-specific network adaptations, with cannabis users showing extended frontoparietal–attentional integration and alcohol users exhibiting stronger occipital–somatomotor engagement. (B) Radar plots summarize the mean DC across Yeo 7 functional networks for each group (healthy controls = purple; cannabis = blue; alcohol = red). All groups engaged dorsal attention and frontoparietal systems, whereas cannabis users showed reduced limbic–default participation and alcohol users showed elevated visual–somatomotor centrality, reflecting opposite topological shifts in predictive connectivity profiles.

In healthy controls, high-DC regions were concentrated in the medial prefrontal and posterior cingulate cortices, orbitofrontal cortex (OFC), and adjacent parietal and occipital areas within default, limbic, and visual networks—fifteen of twenty high-centrality parcels showed direct correspondence with the meta-analytic circuit. These areas likely represent normative components of valuation and contextual processing that distinguish controls from substance users.

In cannabis users, predictive hubs extended beyond this canonical set into the frontoparietal control and dorsal attention systems, with sixteen of twenty high-DC parcels overlapping with drug-related regions. Strong correspondence was observed in bilateral frontal eye field (FEF), parietal, and orbitofrontal territories, suggesting that classification relied on altered integration between attentional and motivational control networks. This pattern aligns with the notion that cannabis-related alterations manifest in the coordination between cognitive control and motivational salience circuits rather than within core valuation areas alone.

In alcohol users, overlap patterns shifted toward occipital, somatomotor, and ventral prefrontal cortices. Fifteen of the twenty predictive hubs coincided with the drug-related meta-analytic map, prominently within visual, sensorimotor, and orbitofrontal regions, indicating enhanced coupling between sensory-driven and interoceptive valuation systems. These findings point to a bottom-up bias in network integration that distinguishes alcohol use from both cannabis use and healthy controls.

This network-level summary was derived from the validated set of top-degree parcels per group, filtered against the full 90-region parcellation to exclude ROIs with missing or low-confidence DC values. At the network level, averaged DC across the Yeo 7 systems revealed consistent involvement of dorsal attention and frontoparietal networks across groups, yet opposite polarity in how predictive influence was distributed: cannabis users exhibited reduced limbic and default participation, whereas alcohol users showed elevated visual–somatomotor centrality (Fig. 3B). Together, these results identify a shared motivational–control backbone that supports classification across groups, while revealing substance-specific shifts in the topology of predictive connectivity—reflecting distinct neural adaptations underlying cannabis and alcohol use.

### Predictive Network Communities and Efficiency Reveal Group-Specific Topologies

To characterize the network organization underlying the discriminative connectivity patterns identified by our classifiers, we applied graph-theoretic community detection to group-level, coefficient-weighted connectivity matrices from each one-vs-rest model. This analysis identified cohesive predictive subnetworks—communities of densely interconnected regions that jointly contributed to group differentiation (Fig. 4).

**Figure 4:**
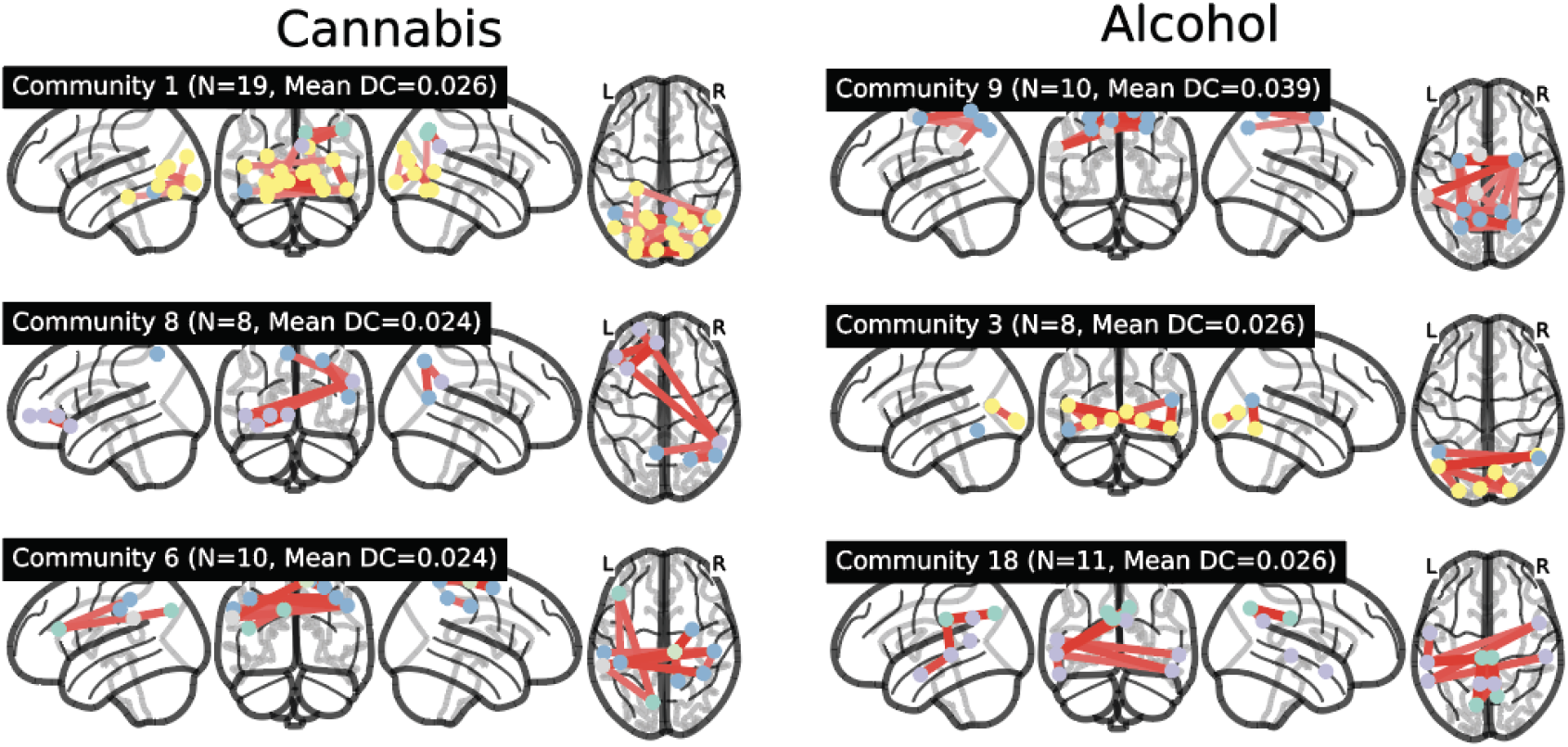
Predictive Community Structure in Alcohol and Healthy Control Models. Predictive network communities derived from the alcohol vs. others (left) and cannabis vs. others (right) classification models. Each row depicts one predictive community, visualized from multiple angles. Nodes represent cortical regions, and edges indicate high-weight predictive connections within each community.

In the *cannabis-vs-others* classifier, three primary communities were identified, encompassing frontoparietal, dorsal attention, and salience regions. Community 1 connected bilateral frontal eye fields, posterior parietal cortices, and medial prefrontal hubs, forming a control–attention circuit. Communities 6 and 8 involved parietal and temporal association cortices, reflecting enhanced coordination between executive and salience systems. Despite preserved large-scale integration, cannabis users showed reduced local efficiency (p < 0.05, FDR-corrected), suggesting decreased functional segregation within these predictive subnetworks.

In contrast, the *alcohol-vs-others* classifier revealed predictive communities anchored in primary visual, somatomotor, orbitofrontal, and anterior insular regions (Communities 3, 9, 18). These subnetworks indicated stronger within-community coupling but reduced long-range integration across control and limbic systems. Alcohol users also exhibited lower local and global efficiency (p < 0.05), consistent with disrupted information flow and a shift toward sensorimotor- and interoceptive-driven processing.

Across both classifiers, overlapping posterior midline regions—including the precuneus and posterior cingulate cortex—formed a shared attentional scaffold, suggesting a common domain-general integration hub. However, the direction of reorganization diverged: cannabis users retained distributed but inefficient integration among frontoparietal and salience systems, whereas alcohol users displayed widespread topological degradation dominated by bottom-up sensory–valuation coupling. These findings indicate that predictive network communities capture substance-specific alterations in mesoscale topology.

## Discussion

In this study, we developed explainable machine learning models capable of accurately classifying chronic cannabis users, alcohol users, and healthy controls based on fMRI-derived functional connectivity. By moving beyond binary classification and demonstrating robust multigroup prediction, we address a critical gap for clinical translation, where distinguishing among multiple substance use profiles is essential. Our approach advances efforts toward biologically interpretable, clinically actionable biomarkers for substance use disorders (SUDs).

A key finding of this work is that connectivity-based models consistently outperformed activity-based models across binary, cross-substance, and three-way classification tasks. This finding extends long-standing views that network-level dysfunctions—alongside localized regional activations identified by prior fMRI studies (Zilverstand et al., 2018; Sutherland et al., 2012)—play a central role in the neurobiology of SUDs. Such network-level frameworks have been extensively discussed in influential models of addiction (e.g., Koob & Volkow, 2016; Lüscher & Voon, 2020), yet these networks have typically been inferred indirectly—from convergent patterns of regional activation and behavior—rather than tested directly. Here, we advance this work by testing network organization in an unbiased manner across substance groups, providing data-driven support for these models. This approach aligns with recent lesion network mapping and machine learning studies that directly test network-level hypotheses in addiction and related disorders (e.g., Stubbs et al., 2023; Joutsa et al., 2022; Siddiqui et al., 2022).

Graph-theoretical analyses further improved model interpretability by identifying predictive network hubs and communities. Regions within the default mode and dorsal attention networks—including the precuneus, posterior cingulate cortex, and frontal eye fields—emerged as central hubs in both healthy controls and cannabis users. These areas are implicated in cognitive control, self-referential processing, and attentional regulation, suggesting that disruption of core control circuits may represent a common substrate of chronic substance use. In contrast, alcohol users exhibited predictive subnetworks centered around somatomotor and salience-related regions, including the insula and orbitofrontal cortex, consistent with known impairments in interoception and habitual control (Goldstein & Volkow, 2011; Naqvi & Bechara, 2009).

Efficiency analyses provided complementary insights into network-level alterations. Cannabis users showed localized reductions in network segregation (local efficiency), whereas alcohol users exhibited broader reductions spanning both local and global efficiency. By demonstrating that multigroup classification is feasible based on connectivity features, our results inherently capture these topological disruptions at the network level.

As noted in a recent review (Ekhtiari et al., 2024), most machine learning studies on addiction have focused on binary comparisons, which may limit their translational potential. This work extends previous approaches by applying three-way classification in a clinically relevant setting. Although the overall multigroup prediction was successful and biologically interpretable, performance differences between cannabis and alcohol users highlight remaining challenges. The lower accuracy for alcohol users may reflect greater neurobiological and behavioral heterogeneity, smaller sample size, or less consistent task-evoked responses.

Future research should include additional substance classes, such as stimulants and opioids, and adopt longitudinal and multimodal designs to improve predictive validity, specificity, and clinical applicability.

In summary, explainable machine learning applied to functional connectivity can distinguish among substance use groups and reveal biologically meaningful neural substrates of addiction, moving the field toward interpretable and clinically useful biomarkers for complex psychiatric conditions.

## Methods

### Participants

We analyzed task-based fMRI data from 166 chronic cannabis users (CU), 136 heavy alcohol users (AU), and 238 healthy controls (HC). All participants completed a cue-induced craving task involving exposure to substance-related and neutral stimuli during scanning.

Cannabis and alcohol user status was determined using self-report assessments and validated screening instruments, including the Alcohol Use Disorders Identification Test (AUDIT) for alcohol users. Participant demographics are summarized in Table 1.

**Table 1.**

|  | Cannabis Users | HC Subjects |
| --- | --- | --- |
| No. of participants | 166 | 124 |
| Sex, Male | 100 (60%) | 53 (43%) |
| Age, Years | 30.1 | 27.4 |
| No. of Drinking Days* | 16.9 (20.38) | 7.48 (14.03) |
| No. of Cannabis Use Days* | 69.02 (15.28) | 0.02 (0.17) |
| *Among Last 90 Days |  |  |
|  | Alcohol Users | HC Subjects |
| Number of participants | 136 | 132 |
| Sex, Male | 106 (77%) | 63 (48%) |
| Age, Years | 28.30 | 29.75 |
| Alcohol Use Disorders Identification Test* | 22.89 | 7.00 |
| *16~19: High risk, 20~40:Addiction likel, |  |  |

### MRI Data Acquisition

Functional MRI data were acquired on a 3T Siemens Prisma scanner using a gradient-echo echo-planar imaging (EPI) sequence (TR = 2000 ms, TE = 27 ms, flip angle = 70°, matrix size = 64 × 64, voxel size = 3.0 mm isotropic). High-resolution T1-weighted anatomical images were also collected for each participant.

### fMRI Data Preprocessing

Functional MRI data were preprocessed using the Nilearn GLM pipeline to ensure consistent quality across datasets. Preprocessing steps included skull stripping, bias field correction, motion correction (MCFLIRT), slice timing correction (AFNI’s 3dTshift), spatial normalization to MNI152 space (ANTs), physiological noise regression (CompCor), and spatial smoothing with a 6 mm full-width at half-maximum (FWHM) Gaussian kernel. Volumes with excessive motion (>3 mm framewise displacement in >5% of volumes) were excluded to minimize motion artifacts. Task effects were regressed out using a general linear model prior to functional connectivity matrix generation, and residual time series were used for subsequent analyses. These procedures replaced earlier SPM-based pipelines, enabling harmonized preprocessing across all participant groups.

### Functional Connectivity Estimation

Following preprocessing, functional connectivity matrices were constructed for each participant. The Schaefer et al. (2018) 100-parcel cortical atlas was used to define regions of interest (ROIs), selected for its balance between anatomical specificity and computational efficiency. Time series were extracted from each ROI, and pairwise Pearson correlations were computed to generate subject-level connectivity matrices for subsequent machine learning analyses.

### Machine Learning Model Training

We trained four types of linear classifiers to predict group membership based on functional connectivity and activity features: L1- and L2-regularized logistic regression, and L1- and L2-regularized linear support vector classifiers (SGDClassifier variants). A One-vs-Rest (OvR) framework was employed to perform multi-class classification among healthy controls (HC), cannabis users (CU), and alcohol users (AU). Hyperparameters, including regularization strength (C for logistic regression; alpha for SGDClassifier), were optimized using a 5-fold cross-validation procedure with grid search on the training data.

Cross-validation performance was evaluated using classification accuracy, area under the receiver operating characteristic curve (AUROC), and F1 scores to ensure model selection balanced sensitivity and specificity. After cross-validation, the best-performing models were retrained on the full training set and evaluated on an independent held-out testing set comprising 20% of participants. Classification performance was assessed using test accuracy, AUROC, and class-wise recall. For imbalanced datasets, such as cannabis users versus healthy controls (∼60/40 distribution), model evaluation emphasized AUROC to mitigate class imbalance effects. In multi-class classification, balanced accuracy (i.e., mean recall across classes) was additionally computed to ensure sensitivity across all groups. In addition to within-substance classification, cross-substance generalization analyses were conducted: models trained to distinguish CU from HC were tested on AU participants, and vice versa. Additionally, to assess connectivity features shared across substance use conditions, we aggregated cannabis and alcohol users into a single substance use group and trained classifiers to distinguish them from healthy controls.

### Three-Class Classification Procedure

Following the binary and aggregated classification analyses, we conducted a three-way classification to simultaneously differentiate healthy controls (HC), cannabis users (CU), and alcohol users (AU). Using a One-vs-Rest (OvR) framework, separate classifiers were trained to distinguish each class against the remaining two. Final class assignments were made based on the highest decision function output, where, for logistic regression models, the decision function reflected predicted probabilities, and for support vector machine models, it corresponded to the signed distance from the decision boundary. Model performance was evaluated using cross-validated accuracy, macro-averaged F1 scores, and area under the receiver operating characteristic curve (AUROC), followed by independent testing on a held-out dataset. Class-wise recall and balanced accuracy were reported to ensure that sensitivity was maintained across all groups, reflecting the practical necessity of accurately identifying multiple substance use profiles in clinical settings.

### Feature Weight Mapping and Graph Construction

Model coefficients from the best-performing classifiers were extracted and multiplied with each participant’s functional connectivity matrix to generate participant-specific weighted connectivity matrices, highlighting predictive inter-regional connections. To focus on the strongest predictive features, weighted matrices were thresholded at 2% sparsity and binarized to construct adjacency graphs.

### Graph-Theoretic Network Analysis

Graph-theoretic analyses were then performed to characterize network-level substrates underlying classification. Degree centrality was computed as the number of connections per node, identifying predictive hubs. Community detection was applied to group-averaged predictive graphs using a modularity maximization algorithm (Louvain method), delineating subnetworks of tightly interconnected regions. To validate the functional relevance of these communities, we repeated the classification using only connectivity features within the identified subnetworks, confirming that they preserved predictive performance. In addition, global and local efficiency metrics were calculated using the Brain Connectivity Toolbox to assess network integration and segregation properties.

### Meta-Analytic Validation of Predictive Hubs

To validate the functional relevance of predictive hubs, top-degree centrality regions were compared against meta-analytic craving-related activation maps obtained from Neurosynth (keywords: “craving,” “drug,” and “alcohol,” thresholded at FDR q < 0.01). Spatial overlaps were quantified for each group and visualized to establish correspondence between predictive network features and known craving-related brain circuits.

**Supplementary Table 1.** Classification accuracy and AUC across models and substance groups.

| Task | Best Model | CV Accuracy | CV Std | Test Accuracy |
| --- | --- | --- | --- | --- |
| 3-Class (Connectivity) | L2 Logistic | 0.808 | 0.036 | 0.783 |
| 3-Class (Activation) | L1 Logistic | 0.501 | 0.059 | 0.52 |
| Binary: Alcohol vs HC (Conn) | L2 Logistic | 0.555 | 0.061 | 0.672 |
| Binary: Alcohol vs HC (Act) | L2 Logistic | 0.443 | 0.068 | 0.509 |
| Binary: Cannabis vs HC (Conn) | L1 Logistic | 0.77 | 0.076 | 0.796 |
| Binary: Cannabis vs HC (Act) | L2 Logistic | 0.602 | 0.07 | 0.373 |
| Aggregated: Users vs HC (Conn) | L2 Logistic | 0.675 | 0.044 | 0.737 |
| Aggregated: Users vs HC (Act) | L2 Logistic | 0.558 | 0.067 | 0.588 |

## Author contributions

A.K. designed the study, integrated analyses and results, interpreted the findings, and wrote the initial manuscript draft. K.R.K. performed the main data analyses, implemented the classification framework, and contributed to figure generation. N.R.D. assisted with machine learning analyses and data preprocessing. M.H. and V.G.F. contributed to data management, pipeline harmonization, and manuscript revision. F.M.F. provided the alcohol and cannabis user datasets and contributed to study design and clinical interpretation. X.G. supervised the project, contributed to study conception and design, and provided critical feedback throughout manuscript preparation. All authors discussed the results, contributed to the final manuscript, and approved its submission.

## Funding and Acknowledgments.

This work was supported in part by the Japan Society for the Promotion of Science (JSPS) through a Grant-in-Aid for JSPS Fellows to A.K. V.G.F. is supported by the National Institute of Mental Health (R21MH129898), and X.G. is supported by the National Institute on Drug Abuse (R01DA043695, R21DA049243). We thank Francesca M. Filbey for providing access to the cannabis and alcohol fMRI datasets, and members of the Center for Computational Psychiatry at the Icahn School of Medicine at Mount Sinai for valuable feedback on data analysis and interpretation.

## References

1. Ekhtiari, H., et al. Functional neuroimaging of substance use disorders: Toward clinical translation. Nat. Rev. Neurosci. 25, 202–220 (2024).

2. Koban, L., Wager, T. D. & Kober, H. A neuromarker for drug and food craving. Nat. Neurosci. 26, 354–365 (2023).

3. Goldstein, R. Z. & Volkow, N. D. Dysfunction of the prefrontal cortex in addiction: Neuroimaging findings and clinical implications. Nat. Rev. Neurosci. 12, 652–669 (2011).

4. Zilverstand, A., Huang, A. S., Alia-Klein, N. & Goldstein, R. Z. Neuroimaging impaired response inhibition and salience attribution in human drug addiction: A systematic review. Neuron 98, 886–903 (2018).

5. Sutherland, M. T., McHugh, M. J., Pariyadath, V. & Stein, E. A. Resting-state functional connectivity in addiction: Lessons learned and a road ahead. NeuroImage 62, 2281–2295 (2012).

6. Koob, G. F. & Volkow, N. D. Neurobiology of addiction: A neurocircuitry analysis. Lancet Psychiatry 3, 760–773 (2016).

7. Lüscher, C. & Voon, V. Revisiting the dopamine hypothesis of addiction: Circuits, plasticity, and behavior. Nat. Neurosci. 23, 564–573 (2020).

8. Naqvi, N. H. & Bechara, A. The insula and drug addiction: An interoceptive view of pleasure, urges, and decision-making. Brain Struct. Funct. 214, 435–450 (2009).

9. Stubbs, W., et al. Lesion network mapping of addiction and compulsive behavior. Nat. Med. 29, 112–123 (2023).

10. Joutsa, J., et al. Human brain networks underlying addiction identified by lesion mapping. Nat. Med. 28, 801–809 (2022).

11. Siddiqui, S. A., et al. Network-based modeling of addiction-related decision-making impairments. Biol. Psychiatry Cogn. Neurosci. Neuroimaging 8, 225–236 (2022).

12. Filbey, F. M., Scharf, M. J., Benschop, A., Daumann, J. & London, E. D. Marijuana craving in the brain. Proc. Natl Acad. Sci. USA 106, 13016–13021 (2009).

13. Filbey, F. M., Yezhuvath, U. S., Scharf, M., Chahal, B. & Kober, H. Cue-elicited neural activity and craving in cannabis dependence. Neuropsychopharmacology 41, 1331–1340 (2016).

14. Filbey, F. M., Claus, E. D., Audette, A. R., Niculescu, M., Banich, M. T. & Brown, S. A. Exposure to the taste of alcohol elicits activation of the mesocorticolimbic neurocircuitry. Neuropsychopharmacology 33, 1391–1401 (2008).

15. Kulkarni, K. R., Kato, A., Fiore, V. G., Gu, X., & Filbey, F. M. Connectivity-based classification of chronic cannabis use with explainable machine learning. Front. Hum. Neurosci. 16, 920345 (2022).

16. Arbabshirani, M. R., Plis, S. M., Sui, J. & Calhoun, V. D. Single subject prediction of brain disorders in neuroimaging: Promises and pitfalls. NeuroImage 145, 137–165 (2017).

17. Dinga, R., Marquand, A. F., Veltman, D. J., Beckmann, C. F., Scheltema, R. A. & Snoek, L. Predicting the naturalistic course of depression from baseline fMRI using machine learning. Biol. Psychiatry 90, 436–445 (2021).

18. Schaefer, A., et al. Local-global parcellation of the human cerebral cortex from intrinsic functional connectivity MRI. Cereb. Cortex 28, 3095–3114 (2018).

19. Tzourio-Mazoyer, N., et al. Automated anatomical labeling of activations in SPM using a macroscopic anatomical parcellation of the MNI MRI single-subject brain. NeuroImage 15, 273–289 (2002).

20. Rubinov, M. & Sporns, O. Complex network measures of brain connectivity: Uses and interpretations. NeuroImage 52, 1059–1069 (2010).

21. Abraham, A., et al. Machine learning for neuroimaging with scikit-learn. Front. Neuroinform. 8, 14 (2014).

22. Esteban, O., et al. fMRIPrep: A robust preprocessing pipeline for functional MRI. Nat. Methods 16, 111–116 (2019).

23. Behzadi, Y., Restom, K., Liau, J. & Liu, T. T. A component based noise correction method (CompCor) for BOLD and perfusion based fMRI. NeuroImage 37, 90–101 (2007).

24. Lancichinetti, A. & Fortunato, S. Community detection algorithms: A comparative analysis. Phys. Rev. E 80, 056117 (2009).

25. Blondel, V. D., Guillaume, J. L., Lambiotte, R. & Lefebvre, E. Fast unfolding of communities in large networks. J. Stat. Mech. P10008 (2008).

26. Yeo, B. T. T., et al. The organization of the human cerebral cortex estimated by intrinsic functional connectivity. J. Neurophysiol. 106, 1125–1165 (2011).

27. Gorgolewski, K. J., et al. Nipype: A flexible, lightweight and extensible neuroimaging data processing framework in Python. Front. Neuroinform. 5, 13 (2011).

